# EEG-based visual deviance detection in freely behaving mice

**DOI:** 10.1101/2021.06.14.448331

**Authors:** Renate Kat, Berry van den Berg, Matthijs JL Perenboom, Maarten Schenke, Arn MJM van den Maagdenberg, Hilgo Bruining, Else A Tolner, Martien JH Kas

## Abstract

The mouse is widely used as an experimental model to study visual processing. To probe how the visual system detects changes in the environment, functional paradigms in freely behaving mice are strongly needed. We developed and validated the first EEG-based method to investigate visual deviance detection in freely behaving mice. Mice with EEG implants were exposed to a visual deviant detection paradigm that involved changes in light intensity as standard and deviant stimuli. By subtracting the standard from the deviant evoked waveform, deviant detection was evident as bi-phasic negativity (starting around 70 ms) in the difference waveform. Additionally, deviance-associated evoked (beta/gamma) and induced (gamma) oscillatory responses were found. We showed that the results were stimulus-independent by applying a “flip-flop” design and the results showed good repeatability in an independent measurement. Together, we put forward a validated, easy-to-use paradigm to measure visual deviance processing in freely behaving mice.

## 1. Introduction

The experiments by Hubel and Wiesel on direction selectivity of neurons in the cat visual cortex (Hubel, 1959; Hubel & Wiesel, 1968) have pioneered a growing scientific field on the visual system and its processing abilities. Since then, the mouse is a widely used animal model to investigate visual processing (Baker, 2013). One important reason is that mice are particularly suitable for genetic modification, such as the use of advanced genetically encoded tools for neuroimaging and neuromodulation that allow unravelling of neuronal network dynamics (Warden et al., 2014). Moreover, transgenic mouse models allow to examine the role of specific cell types or neuronal populations (Sohya et al., 2007; Hamm and Yuste, 2016), as well as to study altered visual processing in the context of human psychiatric disorders (Zhang et al., 2017; Hamm et al., 2020; Perenboom et al., 2020). However, visual processing has hardly been studied in awake, freely behaving mice, as typically head-fixation is used to ensure that visual stimuli reach the eye (Montijn et al., 2016; Carrillo-Reid et al., 2019; Fournier et al., 2020). Assessing measures of visual processing in freely moving mice requires a behavioural setup in which animals are constantly exposed to visual stimuli in their environment irrespective of their bodily position.

Detecting changes in the environment is an important function of sensory systems. The brain can shift attention to changes in the environment via either a passive reduction in the response to redundant stimuli, or an active memory-based increased response to unexpected, or deviant, stimuli (Garrido et al., 2009). The representation of deviance detection in the EEG signal has also been called mismatch negativity (MMN; May et al., 1999). Deficits in deviance detection have been associated with various neuropsychiatric disorders, mainly schizophrenia (Näätänen et al., 2014; Tada et al., 2019). In recent years it has become clear that a homolog of the MMN is also present in the visual modality, the vMMN (Czigler, 2007; Kimura, 2012; Pazo-Alvarez et al., 2003). The vMMN has gained substantially less attention compared to auditory deviance detection and has only twice been studied in rodents (Hamm and Yuste, 2016; Vinken et al., 2017). While these studies were able to assess vMMN, the animals were required to be head-fixated.

Here we set out to develop a novel paradigm to measure deviance-induced differences in visual evoked potentials (VEPs) in freely behaving mice. Based on MMN oddball concepts used in the context of auditory deviance detection (Harms et al., 2016), our vMMN paradigm involves changes in light intensity as standard and deviant stimuli. In order to use the measured EEG waveform difference features for vMMN, the paradigm needs to comply with three principal criteria. First, the paradigm should be able to elicit a *robust deviance response* as measured through the difference between the deviant versus standard VEP responses. Second, the deviance response needs to be *stimulus-independent*, meaning that the same response difference is found when using either of the two stimuli - in our case increases versus decreases in light intensity – as deviant. Third, the VEP deviance effect needs to be repeatable in an independent measurement within the same subject (*repeatability*). After satisfying the three criteria based on VEP waveforms, characteristics of the frequency responses for the paradigm were explored to gain insight in visual deviance-induced oscillatory activity. In addition, the influence of the repeated light stimulation was explored by assessing how the strength of the observed vMMN changed with increasing number of standards preceding a deviant.

## 2. Materials and Methods

### 2.1 Mice

Male C57BL/6J mice (n=13) were used to implement and validate the newly developed vMMN paradigm. Animals were single-housed in individually ventilated cages for at least one week prior to surgeries and maintained on a 12:12 light-dark cycle with *ad libitum* access to food and water. All experiments were approved by the Animal Experiment Ethics Committee of Leiden University Medical Center and were carried out in accordance with ARRIVE guidelines and EU Directive 2010/63/EU for animal experiments. All efforts were made to minimize discomfort of the experimental animals.

### 2.2 EEG implantation surgery

Stereotactic EEG electrode implantation surgery was performed in mice at the age of 2 months. Under isoflurane anaesthesia (1.5%, in oxygen-enriched air), three silver (Ag) ball-tip electrodes were implanted epidurally above the right prefrontal cortex (bregma +2.6 mm anterior, −1.6 mm lateral) and the right and left primary visual cortex (V1; bregma −3.5 mm posterior, +/− 3.0 mm lateral). The relatively lateral V1 position was chosen since multiple studies indicate a role for the visual extra-striate areas (which are located more laterally on the occipital cortex) in the vMMN (reviewed in: Kimura, 2012; Vinken et al., 2017). Two epidural platinum electrodes were placed above cerebellum to serve as reference and a ground electrode, respectively. Electromyogram (EMG) electrodes were placed on top of the neck muscles to record muscle activity. Light-activated bonding primer and dental cement (Kerr optibond / premise flowable, DiaDent Europe, Almere, the Netherlands) were used to attach electrodes to the skull. Post-operative pain relief was achieved by a subcutaneous injection of Carprofen (5 mg/kg). EEG recordings started after a 14-day recovery period.

### 2.3 EEG and VEP recordings

Tethered EEG recordings were performed in a Faraday cage in which animals were connected to the recording hardware via a counterbalanced, low-torque custom-build electrical commutator. Signals were three times pre-amplified, band-pass filtered (0.05 to 500 Hz), then amplified 1200 times and thereafter digitized (Power 1401, Cambridge Electronic Devices, Cambridge, UK) at a sampling rate of 5000 Hz. For the recording of VEPs, mice were placed inside a computer-controlled custom-built LED-illuminated sphere in which tethered mice were able to move freely (Van Diepen et al., 2013). The sphere (30 cm diameter) was coated with high-reflectance paint that spread light produced by a ring of white monochromatic LEDs at the top of the sphere around an opening for the swivel. A baffle prevented the mice from looking directly into the LEDs. After connecting mice to the setup in the sphere, animals were allowed to habituate for at least 10 min. Mice were tested once in a baseline assessment and twice in an oddball paradigm, all on separate days. The baseline assessment, in which a train of light flashes of increasing intensity was presented to the animals, was performed to determine VEP signal quality. 60 flashes of 1 ms with increasing light intensity between ~0.4 to 1.1 μW/cm^2^/nm were presented at 2 Hz, and 5 flashes of increasing intensity between ~1.4 to 2.2 μW/cm^2^/nm at 0.5 Hz. The paradigm was repeated 50 times with 20 s rest in-between blocks.

### 2.4 Visual oddball paradigm

To measure vMMN, a light intensity-based oddball paradigm with decreases and increases in light intensity was developed (Fig. 1). To ensure stable levels of light-adaptation before onset of the oddball sequence, the paradigm started with 10 min of constant light of medium intensity (0.15 μW/cm^2^/nm). Subsequently a 7-minute sequence started in which 300-ms pulses of increased (1.7 μW/cm^2^/nm) or decreased light intensity (0.02 μW/cm^2^/nm) stimuli were interspersed by a 500-ms inter-stimulus interval of the 0.15 μW/cm^2^/nm constant light intensity (Fig. 1). The constant level of light in between the sequence of standard and deviant stimuli was used to prevent occurrence of dark adaptation between stimuli. The intensities of increases and decreases were chosen based on VEP amplitudes in the grand average baseline curve in such a way that the amplitude change from decrease to ISI level was the same as the amplitude change from ISI to increase level. The stimulus duration of 300 ms was based on earlier vMMN studies that used stimulus durations between 80 and 500 ms, in humans (Stagg et al., 2004; Kimura et al., 2010; Sulykos and Czigler, 2014) and rodents (Hamm and Yuste, 2016; Vinken et al., 2017). Deviant stimuli were semi-randomly spread through the sequence, with the constraint of a minimum of two standard presentations before the next deviant. The first stimulation block in the paradigm contained 500 stimuli, 437 (87.4%, the standard) of which were intensity increases and 63 (12.6%, the deviant) of which were light intensity decreases. After this block, the paradigm (including the 10 minutes constant light at the start) was repeated with a swap of standard and deviant stimulus type. This so called ‘flip-flop’ paradigm allowed for assessment of differences between standard and deviant stimuli irrespective of stimulus type (Harms et al., 2016), in our case increased *vs* decreased light intensity. The visual oddball paradigm was performed twice for every animal on separate days. Every animal was once tested in the morning (1^st^ half of the light phase) and once in the afternoon (2^nd^ half of the light phase), whereby the order of the morning and afternoon measurement was counterbalanced over the animals.

**Figure 1.**
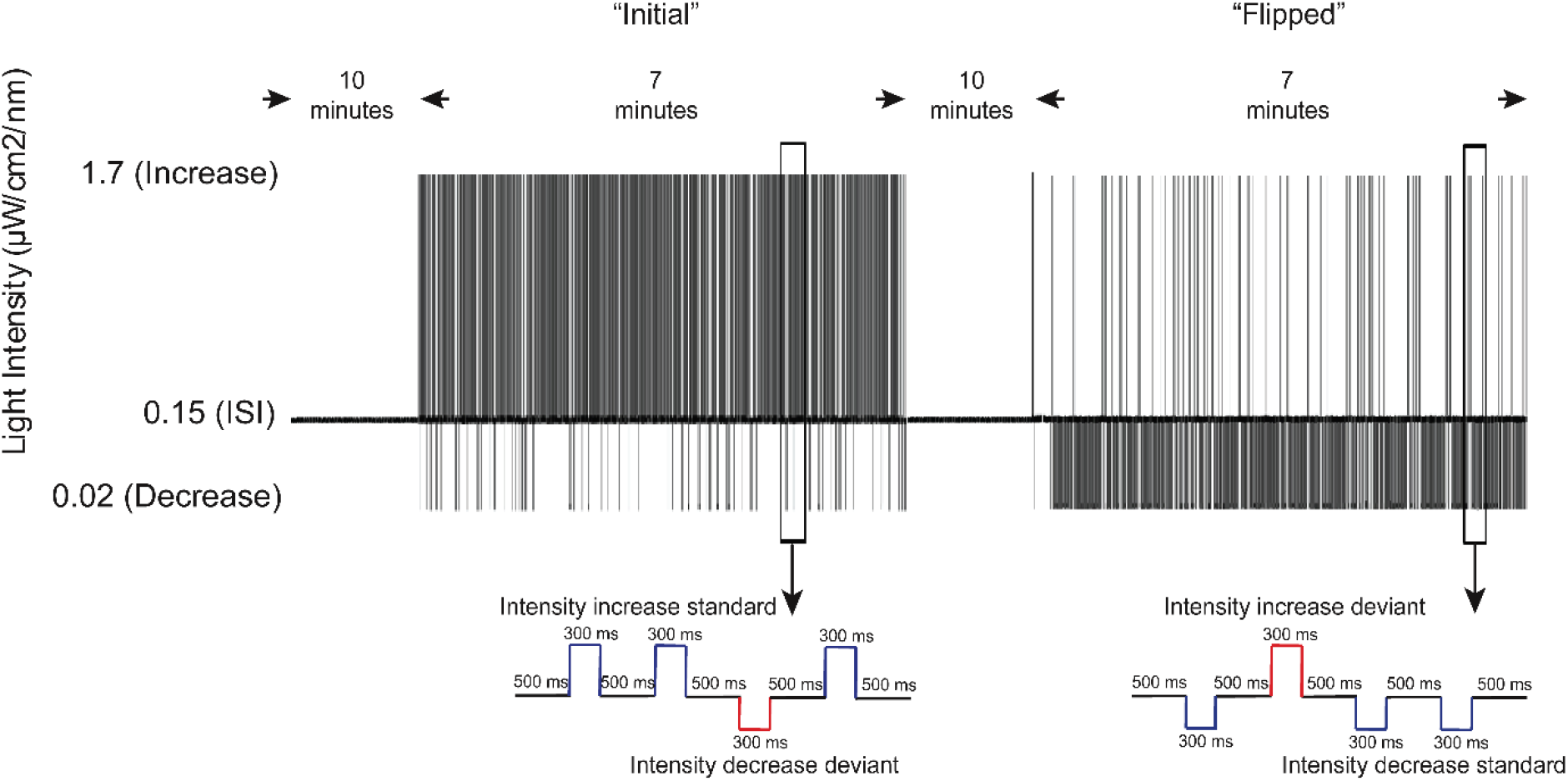
Graphical representation of the light-intensity oddball paradigm used for visual mismatch negativity in freely behaving mice. Mice were presented with an oddball paradigm with increases (1.7 μW/cm^2^/mm) and decreases (0.02 μW/cm^2^/mm) in light intensity as stimuli, with intermittent intermediate intensity levels (0.15 μW/cm^2^/mm). The paradigm was presented as a ‘flip-flop’ in which the “initial” presentation with intensity increase standards and intensity decrease deviants (left), was followed by a “flipped” presentation with intensity decrease standards and intensity increase deviants (right). Initial and flipped stimulation blocks lasted ~ 7 min each. Before the initial stimulation block and in between the initial and flipped stimulation blocks, 10 min of constant intermediate light (0.15 μW/cm^2^/nm) was presented. For the analysis, standards of increased intensity were compared to deviants of increased intensity, and standards of decreased intensity are compared to deviants of decreased intensity.

### 2.5 Analysis

No animals had to be excluded on the basis of low signal quality as judged from the baseline assessment of stimulus responses. For two animals, positive-negative inverted signals were evident on one of the visual cortex electrodes (once right V1 and once left V1); these electrodes were excluded from analysis. Next, recordings were manually checked to exclude recording periods with artefacts, as well as periods of sleep, as deviance detection is known to be attenuated or even absent in non-REM sleep (Sculthorpe et al., 2009). For sleep detection, recordings were first screened for the presence of periods where a passive infrared (PIR) motion detector did not pick up non-specific locomotor activity. If periods without locomotor activity were present, they were checked for the presence of non-REM sleep, as defined by high amplitude delta (<4 Hz) waves, so called slow waves, in the frontal EEG signal in combination with an absence of activity in the EMG signal. Two recordings which contained periods of sleep were excluded from EEG analysis (both being the first recording of the animal). Additionally, seven recordings were excluded from the analysis of locomotor activity, since sleep episodes were present in the baseline periods. For one animal both the first and the second recording were excluded due to the presence of sleep, this animal was thus not included in the locomotor activity analysis.

#### 2.5.1 VEP waveforms

Data pre-processing was performed in Matlab (Versions 2018a & 2018b, MathWorks, Natick, MA, USA). EEG data were low-pass filtered at 70 Hz with a fourth order Butterworth filter. For evoked potential waveform analysis, VEPs were extracted from the data of each recording electrode from 50 ms before until 300 ms after stimulus onset. Subsequently, VEPs were grouped into deviant and standard stimuli, irrespective of being a light intensity increase or a light intensity decrease. Within those two categories, trails were averaged, and subsequently baseline corrected, using a latency window that ranged from −50 to 0 to ms prior to the change in light intensity. For plotting purposes, all 437 standard trials were averaged. However, to have a balanced numbers of standards and deviants in the statistical comparisons, bootstrapping of 100 random sets of 63 standards was used in the statistical analysis. Difference waves were calculated by subtracting the standard from the deviant VEP. A comparison between the difference waves of the right and left V1 electrode (using cluster-based permutation analysis) did not reveal any time windows of significant differences (data not shown). In subsequent analyses VEPs from the right and left electrode were averaged. Averaging over trials, electrodes and recordings was performed for the data from individual animals before performing any statistical analysis for the data-sets across animals.

To test whether vMMN was significantly different from zero cluster-based permutation analysis was used as previously described (Maris and Oostenveld, 2007). In short, dependent *t*-test statistics were obtained for every time point (0.2-ms steps) of the difference waves and were clustered over time along adjacent points that reached above the *t*-value threshold corresponding to an alpha-level of 0.05. The sum of all *t*-values in a cluster was used as the cluster statistic. To assess significance of these clusters, a ‘null’ distribution was created by performing 1000 random permutations with the individual animal difference waves and zero. Cluster statistics were extracted for every permutation in the same manner as described above. Both the largest positive and the largest negative cluster from each permutation were used to create two distributions. Clusters in the actual data were considered significant when exceeding the 97.5-percentile threshold for cluster size in either the positive or negative distribution. The permutation process was repeated for difference waves computed with each of the 100 randomly selected subsets of 63 standards. The largest cluster for each component (the early (30-70 ms), the late (70-150 ms) and effects after vMMN latencies (> 150 ms)) was saved into a p-value distribution of which the median, the maximum, the minimum and the percentage of p-values below alpha (p < 0.05) were reported (median [min max], percentage). Medians were reported instead of means, because the p-values were not normally distributed. When no cluster was found, a value of 1 was added to the distribution.

Comparable procedures were used to compare VEP features between right and left electrodes, light intensity increases and decreases, and first and second recordings. However, in these cases permutations were performed by randomly exchanging the data between the two conditions in the comparison. As small numbers of clusters were found for the comparison of the first and second recording, for these data all clusters that were found in the bootstrap were pulled together; for the deviants, no bootstrap was used but all 63 deviants were simply compared between the first and second recording.

Cluster-based permutation analysis does not have a good level of precision for finding exact on- and off-sets, therefore borders of the time, as well as time-frequency, windows of reported clusters should be interpreted carefully (Sassenhagen and Draschkow, 2019). Latency windows plotted in the VEP figures display the windows as found by the analysis with all standards.

#### 2.5.2 Time-frequency analysis

For analysis of time-frequency responses (TFRs), single trial data (i.e., from a single stimulation; either a standard or deviant) were extracted from the EEG signal from 1 s before to 1.5 s after stimulus onset. The data was low-pass filtered at 150 Hz. Like with the VEP analysis, trials were grouped into standards and deviants irrespective of the stimulus being a light increase or decrease. Using the FieldTrip toolbox for EEG/MEG-analysis (Oostenveld et al., 2011; Donders Institute for Brain, Cognition and Behaviour, Radboud University, the Netherlands), Hanning window convolution was performed with 5-ms time windows on single trials. Frequencies were extracted from 4 - 150 Hz with 1Hz linear steps. The number of cycles increased from 2 to 10 with increasing frequency. Next, power was converted to a log10 scale and an absolute baseline correction was performed using a window from 200 until 100 ms before stimulus onset as the baseline. This window was chosen to avoid including stimulus related activity that would be smeared (in time) due to the width of the Hanning window. The average time-frequency map of standard trials was subtracted from the average time-frequency map of deviant trials. Additionally, to assess non-phase-locked TFRs, per condition the *average* VEP response was subtracted from individual trials in the time domain before performing the same time-frequency analysis as described above (Cohen, 2014; Park et al., 2018). To test for statistical significance of clusters in the TFR difference maps, cluster-based permutation analysis was used as described above for VEP waveform analysis. T-test statistics were in this case obtained for every time-frequency point (5-ms and 1-Hz steps) and clustered over time and frequency. TFR analysis was performed with the full set of standards.

#### 2.5.3 Exploratory analysis of the effect of preceding number of standards

To explore effects of the number of standards since the last deviant, in other words the number of preceding standards, on visual evoked EEG responses (both VEPs and TFRs) including vMMN amplitude, linear mixed effects modelling was performed (Bates et al., 2015a; Kuznetsova et al., 2017). Mixed models have a significant advantage over traditional regression models since they consider the number of individual trials that contribute to a condition, as opposed to calculating the unpooled means per animal, per condition, and losing this type of information (Gelman and Hill, 2007). Here, fewer observations were available for the higher number of trials since the last deviant. Models were estimated and analyzed using the R-package *‘lme4’* (Bates et al., 2015b, RStudio, version 1.2.5042 (R-version 4.0), Boston, MA, USA; lme4 package version 1.1-27; Bates et al., 2015b) and *‘lmerTest’* (Kuznetsova et al., 2017).

The amplitude of the VEP waveforms and power of the TFRs were inspected as a function of the number of trials since the last deviant, for both standards and deviants, and light increases and decreases. The VEP waveform mean amplitudes were extracted from each individual trial in the latency windows that were found to be significant clusters in the evoked potential analysis, resulting in two separate models for an early (40 to 60 ms) and a late (70 to 150 ms) latency window. Similarly, the mean frequency power from each of the TFR clusters (across frequencies and time) that were found to be significant was also extracted. For each measurement type (mV or power) and interval, a linear mixed model was constructed with the following formula:

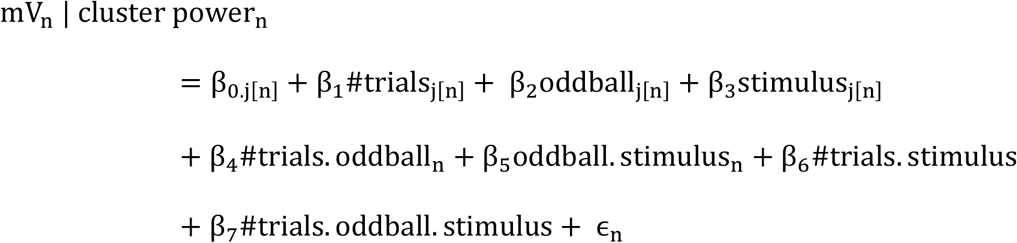

In this formula, for each trial n, the VEP and TFR amplitudes were described by an intercept *β_0_*, a *β_1_* which indicates the number of trials since last deviant (1-30), a *β_2_* which relates to whether the trial was a standard or a deviant and *β_3_* which indicates whether the trial was a light increase or decrease. Finally, *β_4_* – *β_7_* indicates the interactions between number of trials since the last deviant, whether a deviant or standard was presented, and whether the stimulus was a light increase or decrease and ∈_n_ is the residual error term. The subscript j[n] would indicate whether the model included a random effect by animal. To establish the random-effects structure of the model we used a procedure in which we started with a full model (containing random slopes per animal for all corresponding fixed effects; including the interaction terms) and subsequently reduced model complexity by stepwise removing random factors (starting with the higher-order interaction terms) until the model did not improve (AIC selection criteria; (Bates et al., 2015a). This stepwise procedure has been shown to result in a ‘hybrid’ model that avoids overfitting the data while containing relevant random effects (Luke, 2017; Matuschek et al., 2017). As a result, the random effect per subject for the interaction between number of trials since last deviant and whether the trial was a deviant or standard was excluded. T-statistics were used to assess statistical significance of model parameters using the Satterthwaite estimation of degrees of freedom as implemented in the R-package *‘lmerTest’* (Kuznetsova et al., 2017).

#### 2.5.4 Locomotor activity

Locomotor activity of the mice during the vMMN paradigm was assessed by analyzing the activity counts recorded by a PIR movement sensor. Detected movement events were divided into baseline (20 minutes before stimulation and the 10-minute inter-block interval) and stimulation events. Events within stimulation blocks where further subdivided into events during the light increase standard block and the light decrease standard block. For all recordings without sleep episodes, the number of activity counts per minute were calculated per animals, combined over the first and (when applicable) second recording.

Differences in locomotor activity intervals between baseline and stimulation phases, as well as between standard increase and standard decrease blocks were tested with Wilcoxon ranked sum tests, as the data did not pass a one-sample Kolmogorov-Smirnov test for normality.

VEP figures were constructed in GraphPad Prism (Version 8, GraphPad Software, San Diego, CA, USA). Figures of the TFR were constructed in Matlab (Version 2018a,b). Figures of the mixed linear modelling data were constructed in RStudio. Data in text are presented as mean ± standard deviation. The type of variance presented in figures is specified in the figure legends. For all analyses p < 0.05 was considered significant. All data and analysis code (R and Matlab) is available on the OSF data repository (www.osf.io/6bhwf/).

## 3. Results

### 3.1 Visual mismatch negativity can be assessed in freely behaving mice

For the development of the vMMN paradigm for freely behaving mice, we designed an oddball paradigm with sequences of 300-ms white light pulses of increased (1.7 μW/cm^2^/nm) or decreased (0.2 μW/cm^2^/nm) light intensity, interspersed by a 500-ms interstimulus interval at constant light of intermediate intensity (0.15 μW/cm^2^/nm, Fig. 1). Deviant stimuli (63 of 500 stimuli, 12.6%) were semi-randomly spread throughout the sequence with the constraint of a minimum of two standard presentations before the next deviant. In the paradigm both increases and decreases in light intensity were presented once as standard and once as deviant (‘flip-flop’ paradigm; Harms et al., 2016, Fig. 1). The paradigm was presented twice, on separate days. For the first analysis, VEP responses were averaged for, respectively, all standard and deviant stimuli, regardless of being a response to a light increase or light decrease. VEPs recorded from the right and left primary visual cortex (V1), and the first and the second measurement were combined.

Visual inspection of the averaged VEPs revealed a clear distinction between standard and deviant waveforms (Fig. 2A). Both for deviant and standard stimuli, VEPs showed an initial N1 negativity around 30 ms after stimulus onset, followed by a broad positivity between ~50 and ~150 ms. Compared to the response to standard stimuli, the deviant N1 deflection was slightly broadened, while the later broad positivity was of lower amplitude than observed for the standard response. Consequently, the difference wave, computed by subtracting the standard from the deviant response, consisted of a bi-phasic negative component, between ~35 and ~150 ms, with a maximum peak amplitude of - 0.048 ± 0.027 mV (Fig. 2B). Cluster-based permutation analysis revealed two deviance-associated components. The early negative component in the difference wave, ~35-60 ms after stimulus onset, was not significantly different from zero (median *p* = 0.069, [min: 0.024 max: 0.116], percentage < 0.05: 13%), whereas the late negative component, ~70-150 ms after stimulus onset, was (*p* = 0.010, [<0.001 0.052], 99%). Our visual oddball paradigm thus meets the first criterion of *yielding a robust deviance response*, as a significant difference in the response to deviant compared to standard light stimuli could be assessed from VEPs recorded from V1 in freely behaving mice. Compared to the V1 EEG recordings, the oddball paradigm elicited no apparent VEP responses at the prefrontal electrode, nor a distinguishable difference wave (data not shown), indicating specificity of the test paradigm to the visual system.

**Figure 2.**
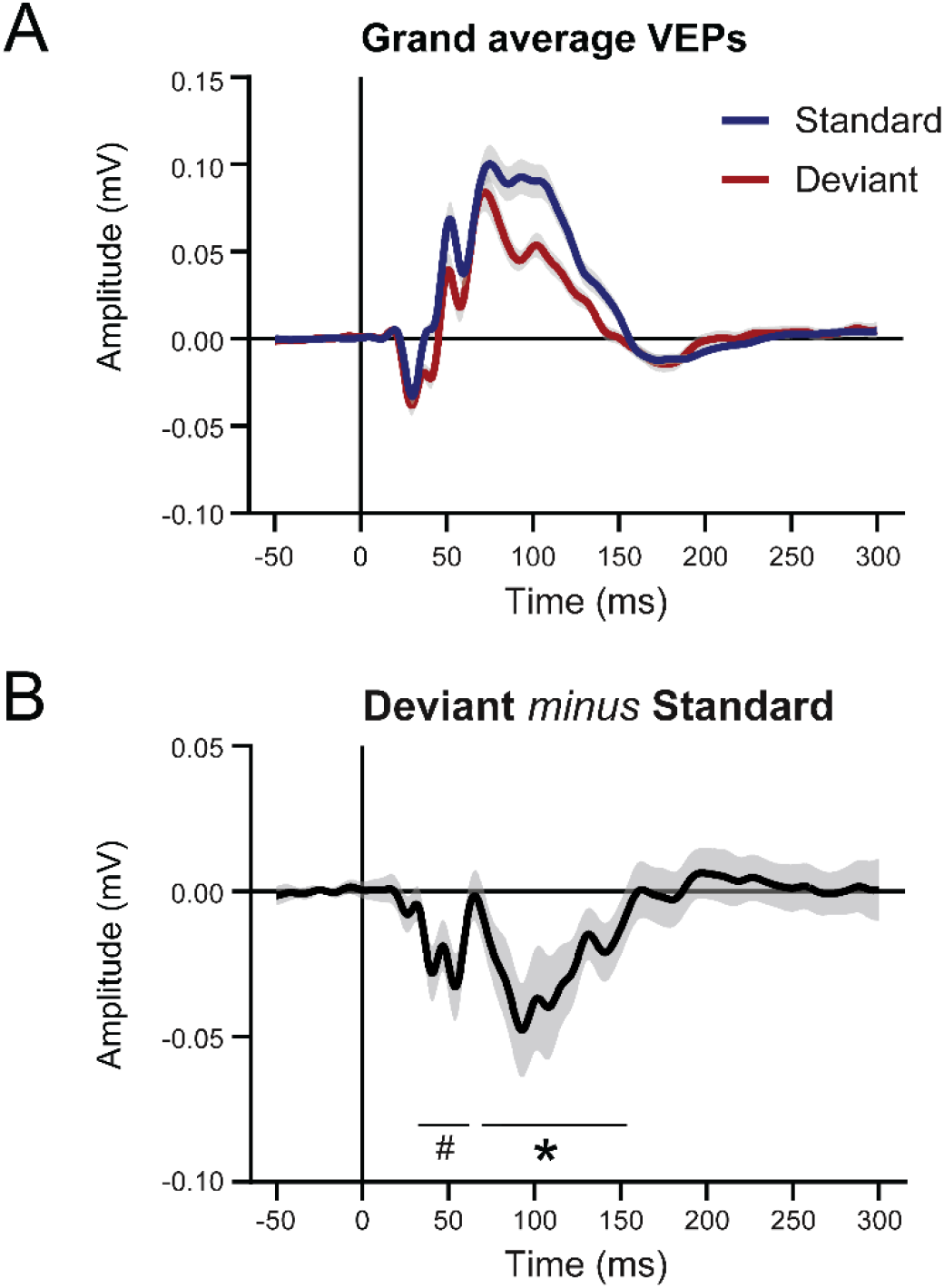
Visual mismatch negativity in the visual evoked potential responses to an intensity oddball paradigm in freely behaving mice. **(A)** Grand average VEP waveforms in response to standard and deviant stimuli. Responses were averaged for, respectively, all standard or all deviant stimuli, independent of the standard or deviant representing a stimulus of increased or decreased light intensity. Responses of the right and left V1, as well as the first and second recording were combined. Data are presented as mean ± standard error of the mean (SEM). (**B)** Deviant minus standard difference wave for the combined ‘intensity increase’ and ‘intensity decrease’ deviants and standards. Data are presented as mean ± 95% confidence interval. Gray shading represents the variance between animals. n = 13, *p<0.01, #p<0.1.

### 3.2 Visual mismatch negativity in the late VEP component is stimulus-independent

To meet the mismatch negativity criterion of stimulus-independency, the difference between VEP responses to standard and deviant stimuli of intensity increases and intensity decreases should contain similar components. Visual inspection of the standard and deviant VEP waveforms (averaged over the responses from V1 left and right, and the two different recording days) revealed different features in the context of intensity increase or decrease stimuli, in particular with respect to the early latencies. Specifically, the VEP in response to an intensity increase, for both standard and deviant stimuli, contained additional early latency components between 20 and 60 ms that were not evident in the VEP in response to an intensity decrease (Fig. 3A).

**Figure 3.**
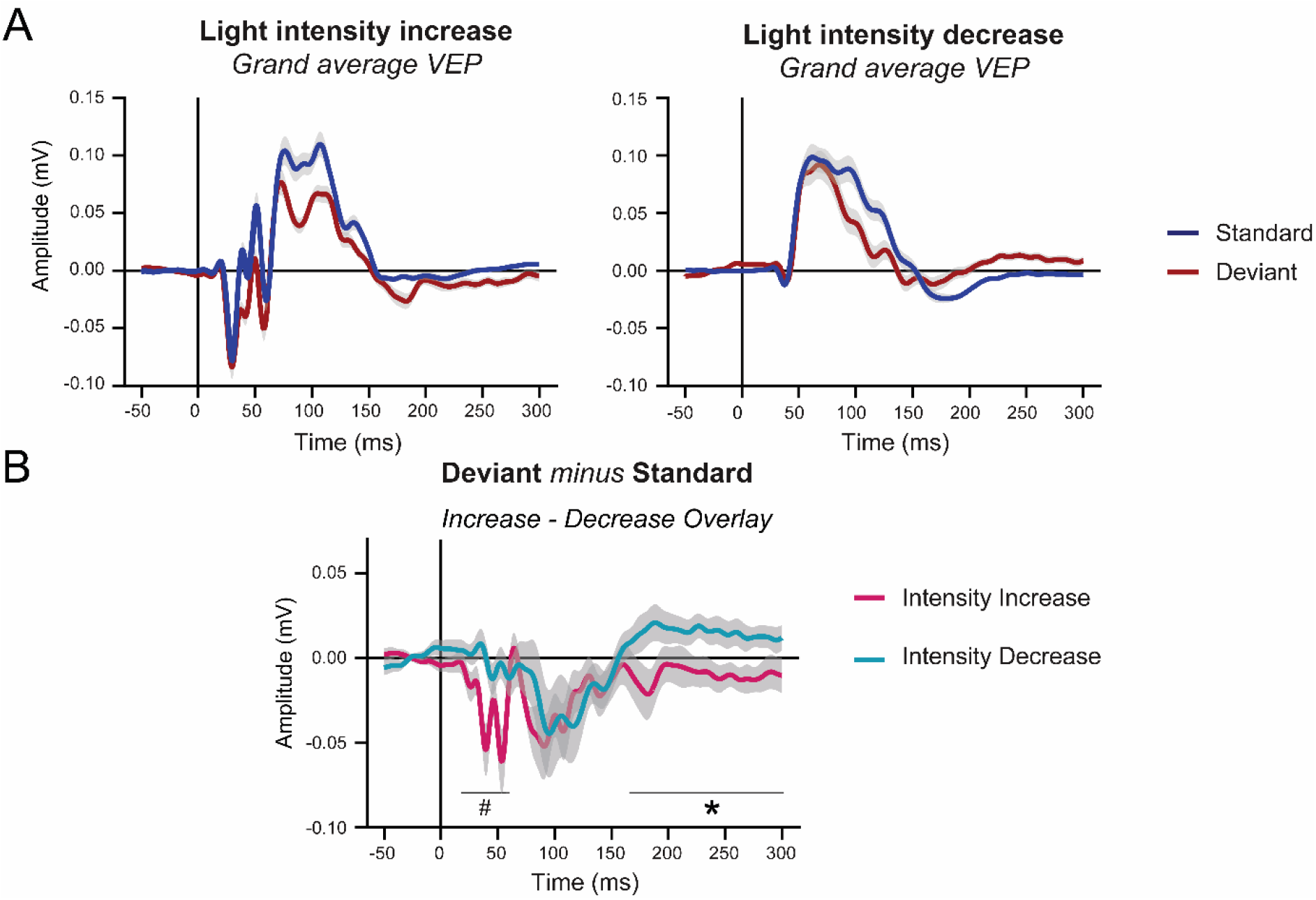
Visual mismatch negativity in the visual evoked potential responses to light pulses of increased or decreased intensity. **(A)** The VEP waveforms for, respectively, ‘intensity increase’ (left) and ‘intensity decrease’ (right) deviants and standards. Data are presented as mean ± standard error of the mean (SEM). (**B)** Overlay of the intensity increase and intensity decrease difference waves. The early negative wave component between 20-60 ms is present only in the difference wave for intensity increase deviants and standards, the late negative wave component around 100 ms is present in both difference waves. A trend level difference between the two difference waves is observed for the early latencies between 20-60 ms. For latencies between 170-300 ms, the waveforms of the intensity increase and decrease difference waves are significantly different. Data are presented as mean ± 95% confidence intervals. Responses were averaged for right and left V1, as well as the first and second recording. Error bars represent the variance between animals. n = 13, *p<0.01, #p<0.1.

While the early components of standard and deviant VEP waveforms for light increases and decreases differed, when subtracting the standard from the deviant response for stimuli of the same light change (i.e. increase or decrease), the deviant-minus-standard difference waves appeared remarkably similar for both light intensity changes with respect to the late component around 100 ms (Fig. 3B). However, the late component of the difference wave, at a latency range of ~70-150 ms, was significantly different from zero for the intensity increase (*p* = 0.006, [<0.001 0.036], 100%), but did not reach significant for the intensity decrease responses (*p* = 0.065, [<0.001 0.242], 42%). The early component of the difference wave was only evident in the difference wave of an intensity increase (p = 0.030, [0.004 0.072], 89%); decrease: *p* = 0.603, [0.106 1.00], 0%). For the difference wave of the intensity decrease responses, the shape of the early component was visible but did not differ in amplitude from zero (*p* = 0.603, [0.106 1.00], 0%). After 150 ms, the difference waves from the intensity increase and decrease responses showed slow shifts in opposite direction which was most evident beyond the ~200 ms latency range of the original VEPs (intensity increase: *p* = 0.149, [0.010 1.00], 25%; intensity decrease: *p* = 0.016, [<0.001 1.00], 65%). When comparing the features of the light increase and the light decrease difference waves directly, the early component was found to differ significantly on a p < 0.1 level (~20-60 ms, *p* = 0.09, [0.012 1.00], 16% smaller than 0.05, 57% smaller than 0.1). Despite the fact that the late component was significantly different from zero for the intensity increase but not the intensity decrease, no differences between the intensity increase and decrease were found for the late component (*p* = 0.368 [0.12 1.00], 0%). In addition, outside the identified window of deviance detection (~30-150 ms), a significant difference between the intensity increase and decrease difference waves was found for the additional late component between ~170-300 ms (*p* = 0.008 [<0.001 1.00], 83%). In conclusion, although both the early and the late latency component were more pronounced in light intensity increase difference waves, the late negative component at ~100 ms could not be statistically differentiated between the responses to light intensity increases and decreases. With the use of this component of the deviant-minus-standard difference waves, our vMMN paradigm thus satisfies our second criterion of *stimulus independency*.

The comparison of the intensity increase and decrease responses also revealed, perhaps not surprisingly, that the ‘off-response’ to an intensity increase – in essence being an intensity decrease – showed a VEP similarly shaped as the ‘on-response’ of the intensity decrease and vice versa (Supplementary Fig. 1). Increases and decreases in light intensity thus seemed to be processed as *shifts* in light intensity rather than as flashes of *different* intensities. The on- and off-responses to a light increase showed slightly higher amplitudes compared to the on- and off-responses to a light decrease. The chosen magnitude of the intensity shifts, which was larger for increases than decreases (i.e. a shift from 0.15 to 1.7 compared to 0.15 to 0.02 μW/cm^2^/mm), was selected based on tests with a 1-ms flash VEP paradigm that showed an equal amplitude difference for both increase and decrease intensities compared to the VEP amplitude response to the ISI intensity. However, in the deviant paradigm the larger intensity shifts still evoked a slightly higher amplitude response. As the latencies of all identified deviance detection components fall within the 300-ms duration of the light stimuli, these off-responses do not affect our deviance detection.

### 3.3 Visual mismatch negativity shows repeatability in an independent measurement

Our third criterion for a vMMN paradigm concerns repeatability of the outcome in independent measurements. To assess this, each animal was subjected to the visual oddball paradigm twice on two separate days. Using cluster-based permutation analysis, no differences between the first and second recording were observed for either the standard VEPs, deviant VEPs or difference waves for the combined responses to intensity increases and decreases (Fig. 4). For both the standard VEP and the difference wave, only four percent of the clusters found after bootstrapping with random subsets of standards was smaller than 0.05, of which all but one had a latency of more than 150 ms (difference wave: *p* = 0.498 [0.010 0.988], 4.1%; standard: p = 0.490 [0.006 0.938], 4.0%). For the deviant, two clusters were found (*p* = 0.814/0.608). These outcomes indicate that our visual oddball paradigm has a good *test-retest reliability* and therefore also meets the third criterion.

**Figure 4.**
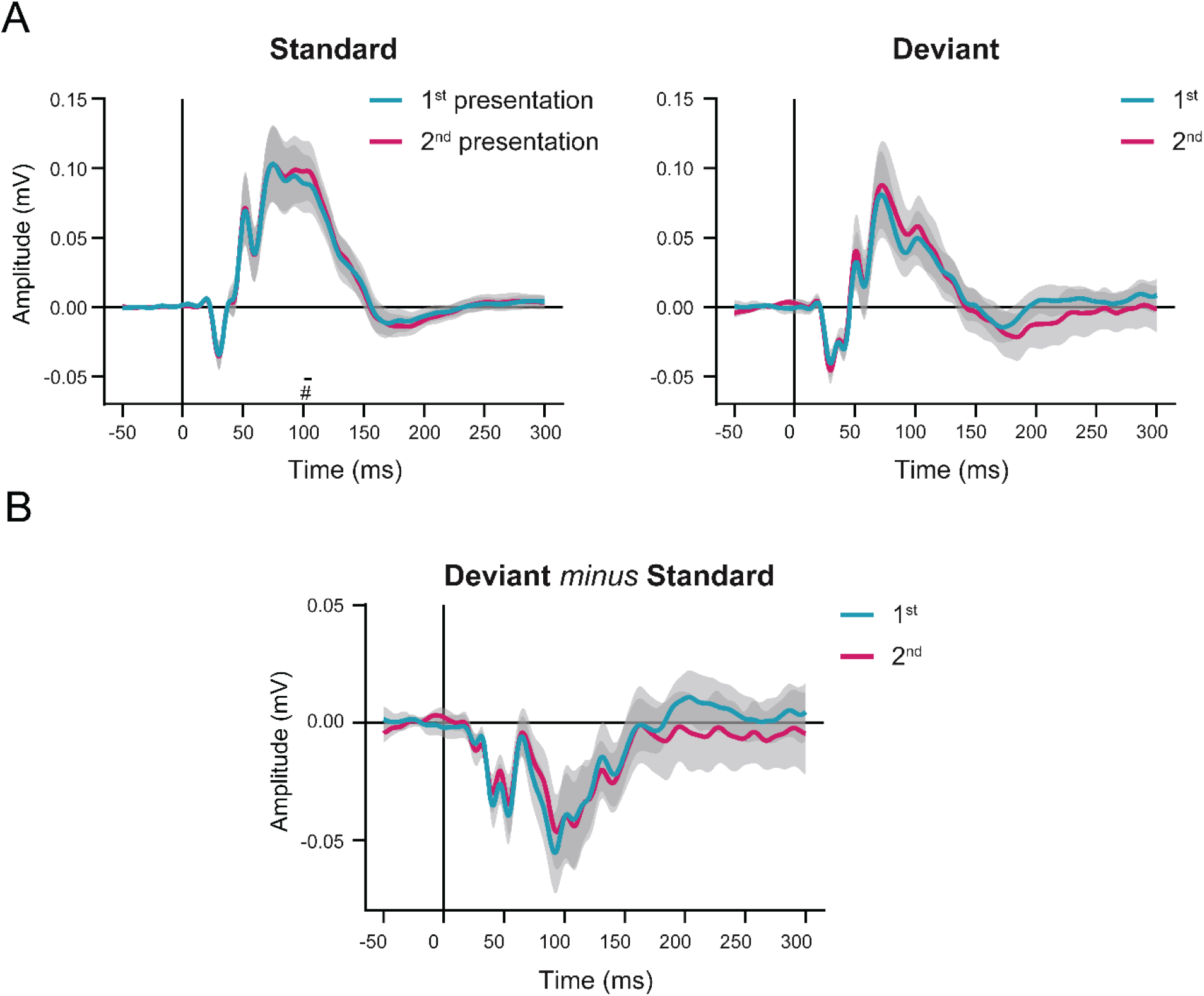
Comparison of the visual evoked potential responses from the 2 independent measurements. The same light intensity oddball paradigm was presented to all mice twice, on separate days (i.e. 1^st^ and 2^nd^ presentation). **(A)** VEPs in response to standard and deviant stimuli, averaged for, respectively, the 1^st^ and the 2^nd^ presentation. **(B)** Overlay of the deviant minus standard difference waves. n = 11, data are presented as mean ± 95% confidence interval. Gray shading represent the variance between animals. Cluster-based permutation analysis did not reveal any significant differences between the 1^st^ and the 2^nd^ presentation.

### 3.4 Mice show intrinsic drive for locomotor activity during visual stimulation

An advantage of using a freely behaving set-up is the opportunity to assess spontaneous behaviour during the EEG recordings. Mice in our vMMN paradigm turned out to show relatively constant high locomotor activity levels during recordings. An exception to this were two mice which were asleep during part of the first recording session, the recordings of which were excluded from further analysis. For the remaining recordings of all mice, periods of inactivity were rare and the overall average interval between two locomotor activity events (as registered by PIR movement sensors in the setup) was only 0.83 ± 0.02 seconds. The occurrence of VEPs during periods inactivity was too infrequent to allow correlating VEP features with expression of locomotor activity. No differences were found between the activity counts per minute for the baseline (72.07 ± 1.80) and the stimulation windows (72.56 ± 2.90, Z = 0.72, *p* = 0.471), nor between the stimulation block in which intensity increases served as standards (73.03 ± 2.52) and the block in which intensity decreases served as standards (72.08 ± 4.76, Z = 0.12, *p* = 0.908). Thus, during the EEG assessments in the light sphere, mice showed a high drive for locomotor activity, which was not affected by the presentation of visual stimuli.

### 3.5 Visual deviance detection is also evident from the light-triggered time-frequency response

In addition to examining VEP waveform features from the deviant-standard difference waves, we analysed the EEG TFR. Human studies showed that vMMN has oscillatory components that are not phase-locked to the stimulus and would therefore cancel out when averaging over trials, which is part of classical event-related potential (ERP) analysis (Stothart and Kazanina, 2013). TFRs are time-locked, but in contrast to ERP waveforms, not necessarily phase-locked to the stimulus and can therefore give a more complete picture of stimulus-associated activity. Visual inspection of the frequency spectra in response to standards and deviants revealed activity in several frequency ranges. The EEG response to standard stimuli – combined for intensity increases and decreases – showed an apparent increased power for the beta-lower gamma (~20-40 Hz, labelled with ‘1’ in Fig. 6A) and the gamma range (~50-100 Hz, labelled with ‘2’ in Fig. 6A) at a latency between ~20 and ~70 ms after stimulus onset. In addition, a broad increase in power was seen for the theta range (~4-9 Hz, labelled with ‘3’ in Fig. 6A), evident from stimulus onset to a latency of ~200 ms. While the TFR to deviant stimuli showed an overall comparable pattern (Fig. 6A), comparison between deviant and standard TRFs in a deviant minus standard heatmap revealed multiple clusters with significantly different frequency components (Fig. 6A). Most evident was a cluster between ~20-120 ms, indicating increased EEG power in the range from ~10-70 Hz in response to deviants (*p* = 0.022, labelled with ‘A’ in Fig. 6A). This cluster seemed to be the result of a combination of an altered shape of the beta/gamma response (labelled with ‘1’) to the deviant compared to the standard stimuli, as well as an additional deviant response in the alpha/beta band (~10-20 Hz, labelled with ‘4’ in Fig. 6A) which was not evident in the response to the standard. The gamma response (~50-100 Hz) contained less power in response to deviant compared to standard stimuli (*p* = 0.048, labelled with ‘B’ in Fig. 6A). Lastly, increased EEG power in the high gamma range (~80-150 Hz) was seen in response to the deviant compared to the standard, both shortly following stimulus onset between ~0-60 ms (*p* = 0.036, labelled with ‘C’ in Fig. 6A) and in a later window between ~90 and ~300 ms (*p* = 0.002, labelled with ‘D’ in Fig. 6A).

**Figure 5.**
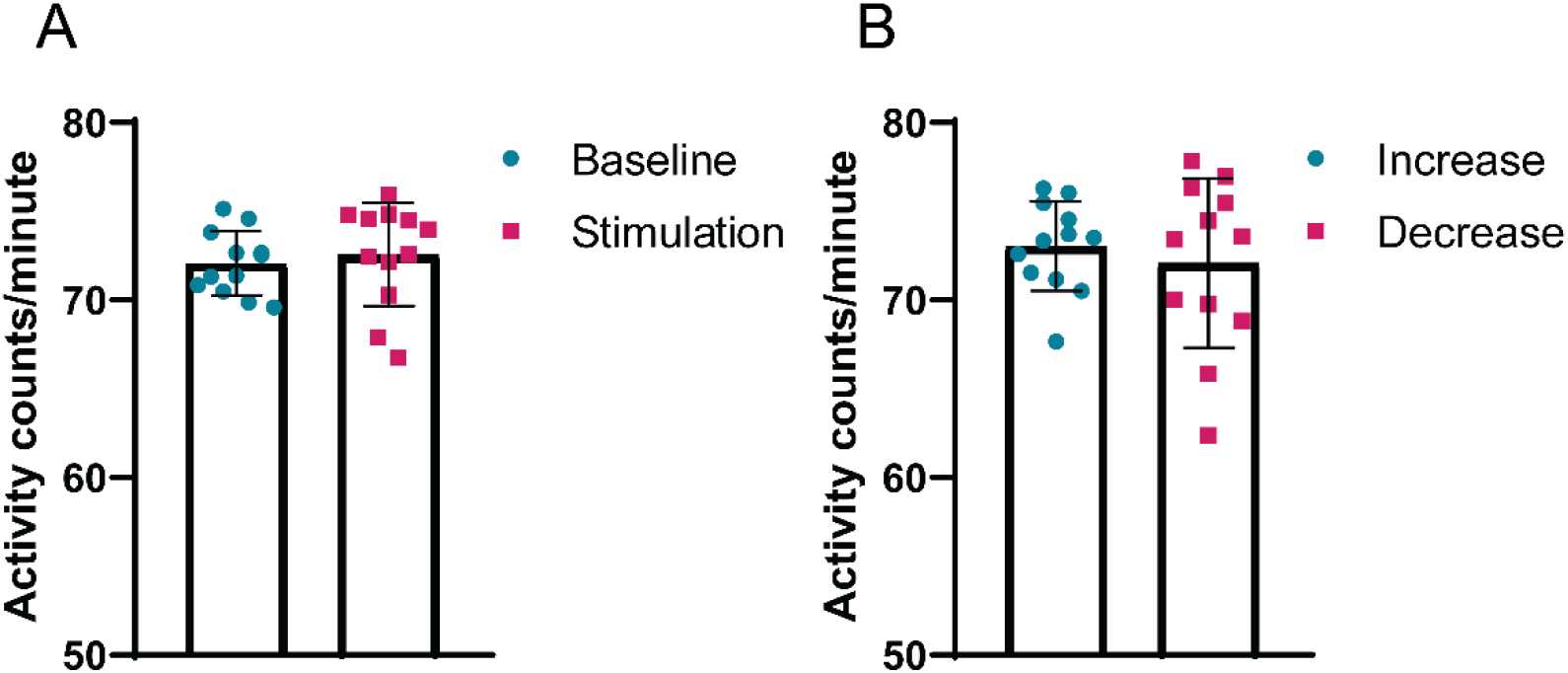
Locomotor activity during EEG assessments in the freely behaving visual stimulation set-up. Non-specific PIR-movement sensors inside the light spheres registered activity-counts. The number of activity-counts per minute were calculated for all animals for the different phases of the paradigm. Locomotor activity was compared between the baseline and stimulation periods **(A)**, and between the stimulation blocks with respectively intensity increase or decreases as standard **(B)**. N = 12, data are presented as mean ± standard deviation with individual data points.

**Figure 6.**
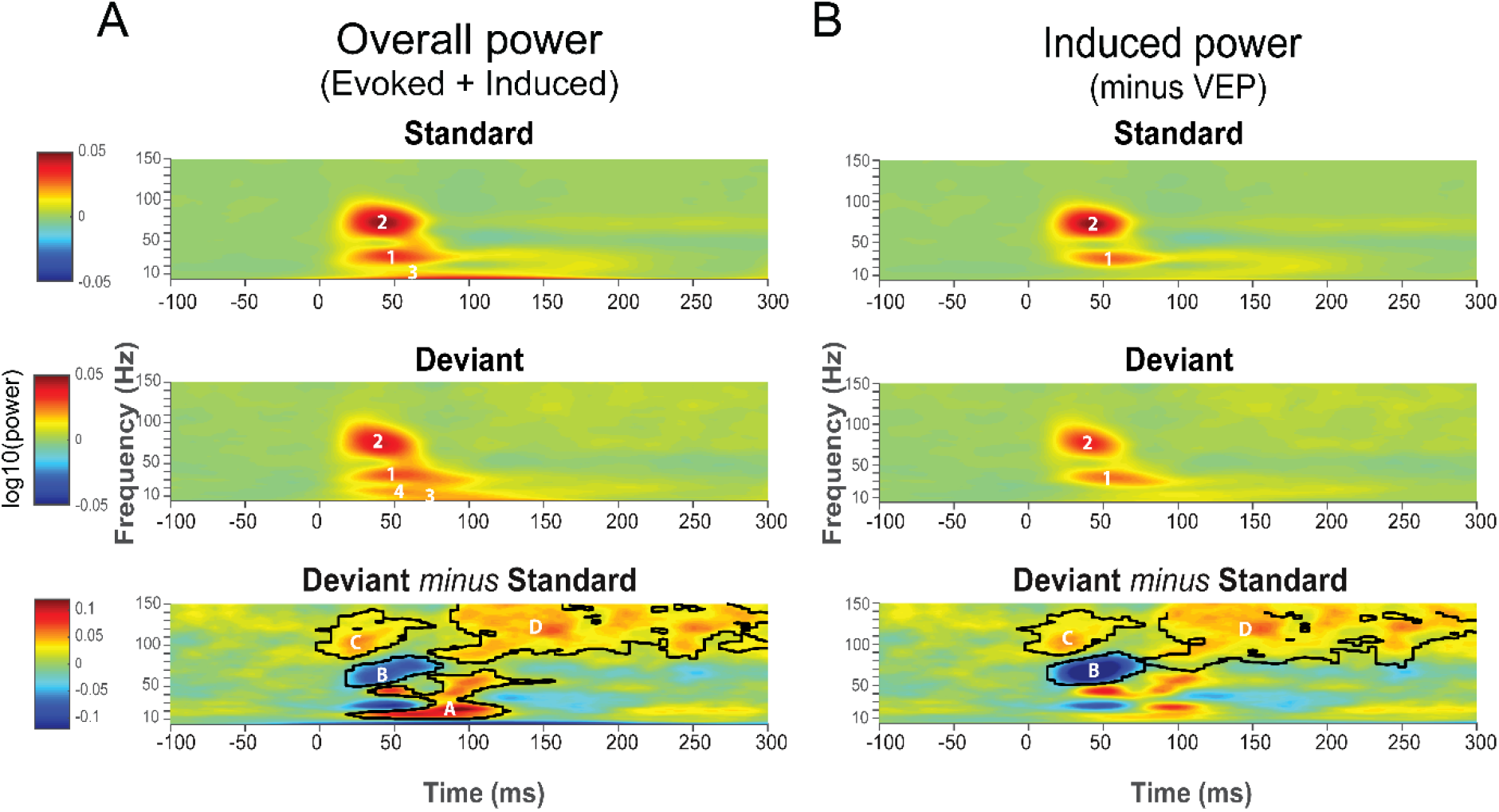
Visual mismatch negativity in the time-frequency response. Panels show clusters of the power of both overall oscillatory activity **(A)**, as well as induced oscillatory activity **(B)** in the vMMN paradigm. To isolate induced oscillatory activity, the averaged waveform was subtracted from each individual trial before running a time-frequency analysis. From top to bottom panel time-frequency responses to standard stimuli, deviant stimuli, and a deviant minus standard difference plot are shown. TFRs were obtained by performing Hanning-window convolution 4-150 Hz with 5 ms time steps. Absolute baseline-correction was performed using −0.2 - −0.1 ms as the baseline. TFRs to light increases and decreases, the right and left V1 as well as and second recording were averaged. Y-axis lower cut-off is 4 Hz. In the difference plot, significant (p<0.05) time-frequency clusters are outlined. n=13.

Oscillatory activity can be divided into evoked power, which is the direct frequency representation of the VEP waveform response, and induced power, which is the oscillatory activity that is non-phase-locked to the stimulus and thus not found in the VEP waveforms (Jones, 2016). To asses which oscillatory clusters in our analysis represented induced power, the time-frequency analysis was also ran after subtracting the average VEP waveform from every single trial per condition (Park et al., 2018). Clusters 3 and 4 in the TFR and cluster A in the difference plot were not present when using this analysis (Fig. 6B) and thus represent evoked power. On the other hand, clusters 1 and 2 in the TFR and clusters B, C and D in the difference plot were still present after running time-frequency analysis on mean-subtracted data and represent the power of induced oscillatory activity. With our visual oddball paradigm in freely behaving mice, deviance detection was thus not only reflected in the VEP waveforms, but also in evoked as well as induced power in the EEG time-frequency responses.

### 3.6 Higher numbers of standards preceding a deviant strengthen visual mismatch negativity

In light of the ongoing debate about the role of adaptation to the repeatedly presented standards in deviance detection paradigms (Garrido et al., 2009; Grimm et al., 2016), we assessed whether stimulus history influenced our VEP-based vMMN. Using linear Mixed effects models (Bates et al., 2015b; Gelman and Hill, 2007) we explored whether the vMMN amplitude (VEP- and TFR-based) changed with varying numbers of standards preceding the deviant. In addition, we assessed whether this was potentially also affected by the stimulus types, i.e., an intensity increase or decrease. The mean amplitude and oscillatory power of standard and deviant VEPs and TFRs were extracted from each trial, for both the cluster-based defined early (40-60 ms) and late (70-150 ms) latency windows rounded to the nearest ten, and the identified gamma (B, ~50-100 Hz) and high gamma (C and D, ~80-150 Hz) induced frequency clusters. We subsequently analysed the amplitude or power using a linear mixed model with the factors ‘oddball’ (standard, deviant), ‘stimulus’ (light increase, decrease), and ‘number of trials since last deviant’ (1-30; continuous and linear variable).

#### Early negativity (40-60 ms)

amplitudes of the vMMN (deviant minus standard) in the early latency increased with the number of trials since the last deviant (1.684*10^-3^ mV (SE: 0.338*10^-3^) per trial). In other words we observed an increase in the early latency vMMN amplitude with longer stretches of standard trials (oddball × # of trials since last deviance: t(25956) = 4.98, *p* < 0.001, Fig. 7A). The amplitude in response to standard VEPs increased with number of trials since last deviant (0.775*10^-3^ mV (SE: 0.176*10^-3^) per trial; t(14.9) = 4.41, *p* < 0.001) and the amplitude in response to deviant VEPs decreased (−0.909*10^-3^ mV (SE: 0.342*10^-3^) per trial; t(211.9) = −2.66; *p* = 0.008). Furthermore, we confirmed our earlier observation that the amplitudes in the early latency window, irrespective of stimulus history, were stimulus specific (oddball × stimulus type: t(25956) = 4.72, *p* < 0.001; light increase: deviant minus standard: −0.0368 mV, SE = 4.35*10^-3^; t(20.6) = 8.46, *p* < 0.001; light decrease: deviant minus standard: - 0.0039 mV, SE = 4.35*10-^3^; t(20.7) = 0.898, *p* > 0.380).

**Figure 7.**
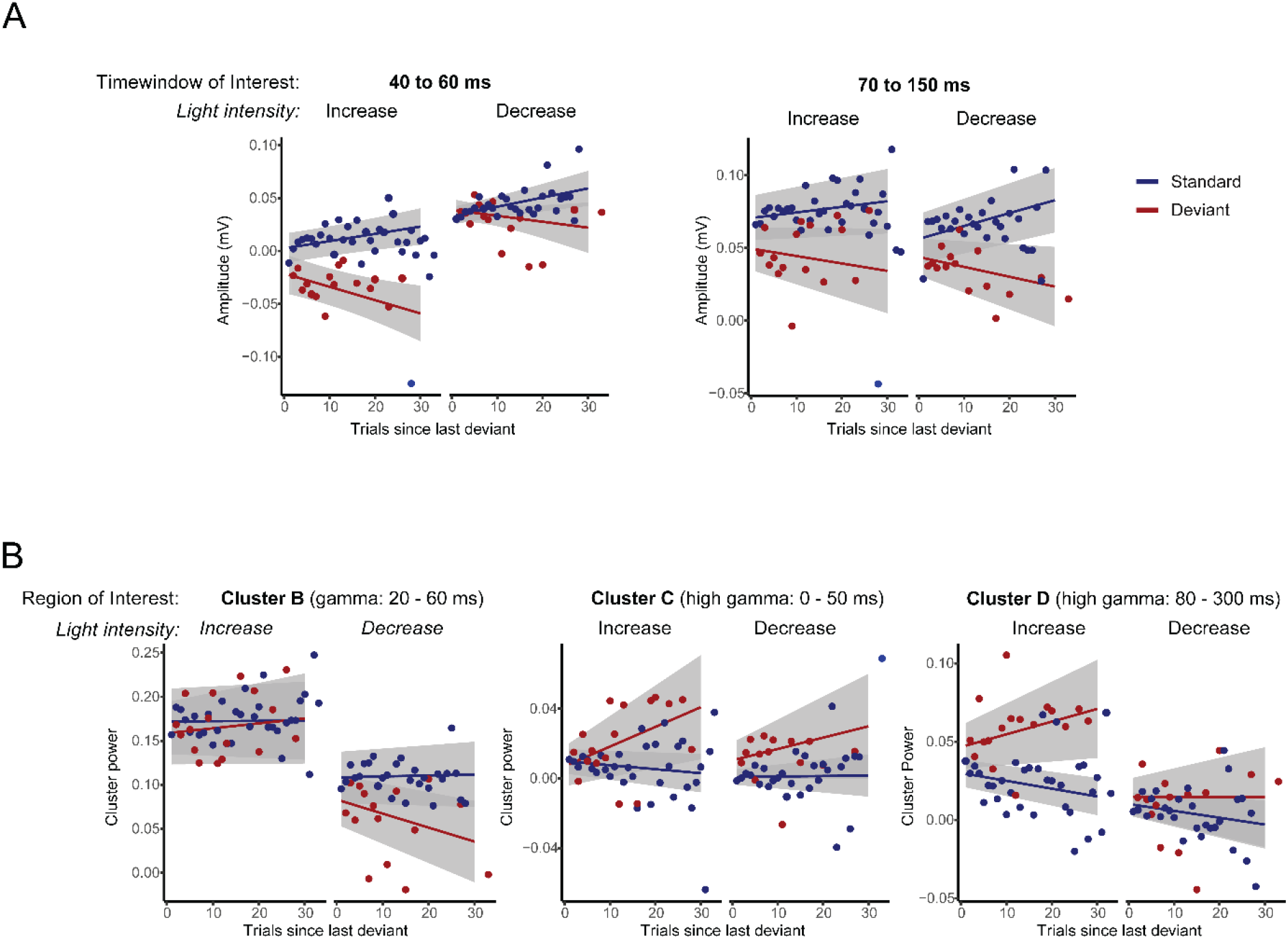
Exploration of effects of stimulus history on VEP- and TFR-based mismatch negativity features. Analyses were performed for the earlier found VEP **(A**, separated for the early and late negativity) and TFR **(B)** components. Each of the different graphs depicts the mean amplitude (for VEP features) or power (for TFR features) as a function of the number of preceding standards since the occurrence of the last deviant. For both VEP and TFR features, the vMMN amplitude (for VEP) or power (for TFR) is the difference between the standard and deviant amplitude, which for the early and late VEP component and TFR cluster B was found to increase with an increasing number of preceding standards. Data are presented separately for standard and deviant stimuli, as well as for intensity increases and decreases. n = 13, data are presented as mean ± 95% confidence interval.

#### Late negativity (70-150 ms)

amplitudes of the vMMN in the late latency window paralleled the observations for the early component with respect to a modulation of the vMMN by stimulus history (oddball × trials since last deviant: (1.232 mV (SE: 0.390*10^-3^); per trial; t(25944) = −3.16, *p* = 0.002). While the amplitude of the late negativity response to standard VEPs increased with decreasing novelty (6.370*10^-3^ mV (SE: 0.180*10^-3^) per trial; t(14.9) = 3.53, *p* = 0.003), for the deviant responses, the late negativity amplitude did not show a significant decrease in amplitude (0.060*10^-3^ mV (SE: 0.383*10^-3^); t(284.3 = 1.55, *p* > 0.122), contrary to what was observed for the early negativity. In the late latency window no overall effect of stimulus type on deviant processing was observed (oddball × stimulus type: t(25944) = −1.19, *p* = 0.235). To further address the stimulus independency, it was tested whether the effects of stimulus history on the late component differed between the stimulus types (i.e. light increase vs light decrease). Addition of this interaction to the model did not improve the fit of the model (ΔAIC = +5.4; ΔBIC = +28, a smaller AIC indicating improvement of the model weighting complexity [number of parameters] and explained variance), supporting the claim that the amplitudes in the late component were indeed stimulus non-specific.

#### Oscillatory clusters

for the TFR-based analysis only the early high gamma cluster (C, 80-150 Hz) showed a significant relationship between stimulus history and deviance detection (C: oddball × trials since last deviant t(25977) = 2.05, *p* = 0.041; D: oddball × trials since last deviant: t(25959) = 1.72, *p* = 0.086, Fig. 7B). However, the large variation and p-values close to the significance threshold suggest that these analyses were underpowered and therefore should be interpreted carefully.

## 4. Discussion

Mismatch negativity is an important function of the brain to identify environmental changes that may require subsequent appropriate behavioural and/or physiological responses. The goal of this study was to develop and validate a method for assessing vMMN in freely behaving mice. The developed paradigm met all three pre-defined criteria: *a robust deviance response, stimulus-independence*, and *repeatability. First*, the light intensity-based oddball paradigm evoked a bi-phasic negativity in the VEP difference wave, of which the late 70-150 ms component was significantly different from zero, indicating that the paradigm was able to assess the ability of mice to differentiate between standard and deviant flashing light stimuli. *Second*, vMMN in this late component was found to be independent of the type of stimulus (i.e., light increase or decrease) that was used a deviant. *Third*, the paradigm showed good repeatability in a second recording performed on a separate day.

The vMMN presented with our paradigm matches well with previously reported vMMN features from both human and rodent EEG. The only other EEG-based vMMN study in mice, in which a pattern-based oddball paradigm was used in head-fixed animals, also showed bi-phasic responses (Hamm and Yuste, 2016). They identified the differences in response to standard and deviant stimuli in early latencies to reflect stimulus-specific adaptation, while differences in later latencies reflected deviance detection activity. Also human visual ERP studies indicated that early components of sensory processing represented adaptation effects, while later components were specifically associated with violations of expectation (Czigler et al., 2006; File et al., 2017). The onset and timing of the early and late phases differed for each of the studies, as well as the present study, both between and within species. Besides differences in neuronal pathways between species, deviance detection latencies may also be influenced by the stimulus complexity (Kojouharova et al., 2019). For example, Hamm and Yuste (2016), which used visual pattern stimuli instead of the light flashes used in our study, found longer latencies (between ~ 40 and 240 ms) in their mouse visual deviance detection features compared to the latencies observed in our paradigm. In our freely-moving deviance detection paradigm, we could not assess the contribution of stimulus-specific adaptation, as the ‘many standards control paradigm’ (Czigler et al., 2006; Hamm and Yuste, 2016; Harms et al., 2016; File et al., 2017) was not used. We were however able to show that the early component was sensitive to stimulus properties (i.e. a significant deviance detection effect was only present for light intensity increases), while the late component was stimulus-type-independent. Additionally, the effect of stimulus history on vMMN amplitude was larger for the early compared to the late component. The finding of larger effects of stimulus history and stimulus-specificity on the early component are in line with the previously reported stimulus-specific adaptation in the early latencies and deviance detection in the later latencies (Czigler et al., 2006; File et al., 2017; Hamm and Yuste, 2016). We therefore speculate that also in our paradigm the early component is primarily driven by stimulus-specific adaptation, while the late component represents active deviance detection. Taken together, our vMMN matches well with that observed in humans as well as in mice. Our data show that head fixation is not required for measuring vMMN in mice, and that the implemented paradigm and observed responses in mice have translational value.

Performing sensory processing assessments in freely behaving animals does not only reduce stress effects but also increases behavioural relevance and can potentially provide more insight via inclusion of behavioural data. In the current study, locomotor activity levels of the experimental animals were recorded during the VEP assessments. This revealed that during the experiments in the light sphere, mice show a high intrinsic drive for locomotor activity as they showed high levels of activity during the entire recording session. Locomotor activity levels were not altered by the presentation of visual stimuli. Since inactive periods were rare, we could not perform separate analysis of VEPs and related deviant features during active and inactive periods. It has previously been shown that locomotor activity affects EEG in rodents (Hansen et al., 2019), however it is unlikely that locomotion contributed to the difference we found between the shape of increase and decrease VEPs since the amount of locomotor activity was equal between both light increase and decrease stimulation blocks. Increasing habituation periods in the set-up, or increasing the duration of the vMMN paradigm might allow for a comparison between VEP and deviant features during active and inactive periods in future experiments. While assessing visual evoked brain activity during full-field visual stimulation is in our view a logic first step, a future goal would be to develop more naturalistic set-ups where ecologically relevant visual input is presented in such a way that the perception of such input is dependent on the animal’s behaviour.

VEPs in response to intensity increases and decreases showed different features in particular for the early components. Rather than being processed as flashes of different intensities, based on the off-responses, the used stimuli seemed to be processed as shifts in light intensity. While the processing of different levels of light intensity (Lopez et al., 2002; Perenboom et al., 2020) and its dependence on light-adaptation (Suzuki et al., 1972) have been studied in mice and cats, little is known on how shifts in light intensity are processed. One early study described the presence of off-responses, which were, contrary to what we observed, of similar shape as on-responses, when light flashes (i.e. increases in light intensity) lasted more than 100 ms (Crescitelli and Gardner, 1961).

Exploratory analysis of the effect of stimulus history showed that an increased number of trials since the last deviant, in other words a longer stretch of preceding standard presentations, increased the amplitude of the vMMN. This was the result of an increased amplitude of standard VEP responses and a decreased amplitude of deviant VEP responses with a higher number of preceding standards. These changes seem to suggest that our VEP-based deviance detection paradigm was sensitive to short-term novelty of the deviant; as it showed larger responses when the previous deviant was presented a longer time ago. The observed positive relationship between vMMN amplitude and number of preceding standards could be a result of stimulus-specific adaptation of the standard, in our paradigm leading to increased amplitudes after more repetitions. It was slightly unexpected that adaptation to the standard after stimulus repetition presented as an increase in amplitude, instead of depression of the responses. This could be interpreted as a sensitization or tuning of the visual cortices to a specific stimulus as shown by others (Clapp et al., 2006; Solomon and Kohn, 2014). While depression-dominated adaptation to visual stimuli has been shown in anaesthetized animals (Keller and Martin, 2015; Sanchez-Vives et al., 2000), a recent study showed that in awake mice certain types of cortical interneurons show depression-dominated while others show sensitization-dominated adaptation, the strength of which changed with locomotion. As a result, the pyramidal responses could be sensitized as well as depressed (Heintz et al., 2020). Alternatively, larger deviance detection after more preceding standards could result from the brain’s response to the violation of a stronger memory-based expectation of the standard (Garrido et al., 2009). Although counterintuitive, violation-alerting activity in our data would actually be represented by the observed reduction in deviant amplitude, resulting in an increased difference with a standard. Further studies are needed to determine which of these two processes primarily drives the deviance detection features in our paradigm. Larger differences in responses to standard and deviant visual pattern stimuli with more preceding standards have previously also been shown in rats, although this difference was dominantly driven by alterations in the responses to the deviant stimuli, without a change in responses to the standard (Vinken et al., 2017). Also in human auditory MMN paradigms the amplitude of the MMN increases when the overall probability of deviants is decreased from 30% to 10% or from 13% to 1.5% (Sato et al., 2000; Sabri and Campbell, 2001), as well as with a higher number of standards preceding a deviant within a paradigm with a stable overall deviant probability of 20% (Matuoka et al., 2006). Together, these findings suggest that the effect size of our vMMN paradigm could be further increased by having a higher minimum number of standards between deviants than the two currently used in our experiments.

In addition to the VEP waveforms, visual deviance detection was also found to be represented in both evoked and induced oscillatory activity. Human visual and mouse auditory studies have previously shown oscillatory responses related to deviance detection, but the paradigms and corresponding responses showed large variability (Stothart and Kazanina, 2013; Ahnaou et al., 2017; Hesse et al., 2017; Yan et al., 2017). Differences across paradigms and species, as well as the fact that some of these studies use auditory while others use visual stimuli, make a direct comparison of findings from the studies assessing frequency response in deviance detection paradigms difficult.

In the TFR the higher gamma clusters between 50-150 Hz represented induced (i.e. non-phase-locked) oscillatory activity. This is in line with the fact that induced power, thought to represent top-down connections, concerns higher frequencies over longer latencies, while evoked power of the visual evoked responses, thought to represent bottom-up connections, concerns lower frequencies over shorter latencies (Chen et al., 2012). The broad increase in high gamma power (80-150 Hz) showed a tendency to be enhanced with more preceding standards, although this effect was not statistically significant Although the role of the various frequency bands in specific functional processes is not well understood, gamma frequency cortical activity has generally been linked to increased spiking activity and network excitation (Yizhar et al., 2011; Cho et al., 2015; Vogt et al,. 2015). In the visual cortex of freely behaving mice, 30-100 Hz broadband gamma activity was found to functionally discriminate between segregated cortical layers of visual processing (Senzai et al., 2019). It was shown that gamma activity can be subdivided into functionally distinct broad- (30-90 Hz) and narrowband (60 Hz) gamma oscillations, which show complementary responses to changes in visual contrast (Saleem et al., 2017). While narrowband gamma has been associated with thalamocortical communication, broadband gamma power is thought to represent corticocortical communication. Our recordings did not allow to distinguish between underlying network mechanisms, but the broad increase in high gamma band activity we observed in the TFR deviant minus standard difference plots could reflect increased corticocortical network activity during deviance processing. This could suggest involvement of the prefrontal cortex, in line with what was found in human vMMN studies (Yucel et al., 2007; Kimura et al., 2010; Kimura et al., 2011), although no robust visual evoked responses were recorded from our prefrontal cortex electrode. The presence of induced broadband gamma responses thus seems to suggest communication between the visual cortex and other cortical, possibly frontal, areas during visual deviance detection. However, the functional significance of EEG activity in certain frequency bands remains to be assessed, thus interpretations related to these specific frequency bands should be drawn carefully.

In conclusion, we developed the first, robust and repeatable vMMN paradigm based on changes in light intensity in freely behaving mice. Our paradigm provides a functional outcome measure for visual processing in these mice. Because no head fixation is needed, our paradigm minimizes animal discomfort while increasing behavioural relevance. The paradigm can easily be implemented to assess sensory processing deficits in mouse models of brain disease, and has the possibility to be compared with experiments in humans which increases translatability of preclinical outcomes.

## Abbreviations

ERP: event related potential
MMN: mismatch negativity
TFR: time-frequency reponse
V1: primary visual cortex
VEP: visual evoked potential

## Acknowledgements

This work was supported by a ZonMW TOP [grant number 91216021, 2017, awarded to HB and MJK] and the national Medical NeuroDelta [awarded to AvdM].

## Author contribution statement

**Renate Kat:** Conceptualization, methodology, investigation, formal analysis, visualization, writing – original draft **Berry van der Berg:** Formal analysis, visualization, writing – reviewing & editing **Matthijs JL Perenboom:** Methodology, software **Maarten Schenke:** Investigation **Arn MJM van den Maagdenberg:** Resources, funding acquisition, writing – review & editing **Hilgo Bruining:** Funding acquisition, writing – review & editting **Else A Tolner:** Conceptualization, writing – reviewing & editing, supervision **Martien JH Kas:** Conceptualization, funding acquisition, writing – reviewing & editing, supervision

## Supplementary figures

**Figure S1.**
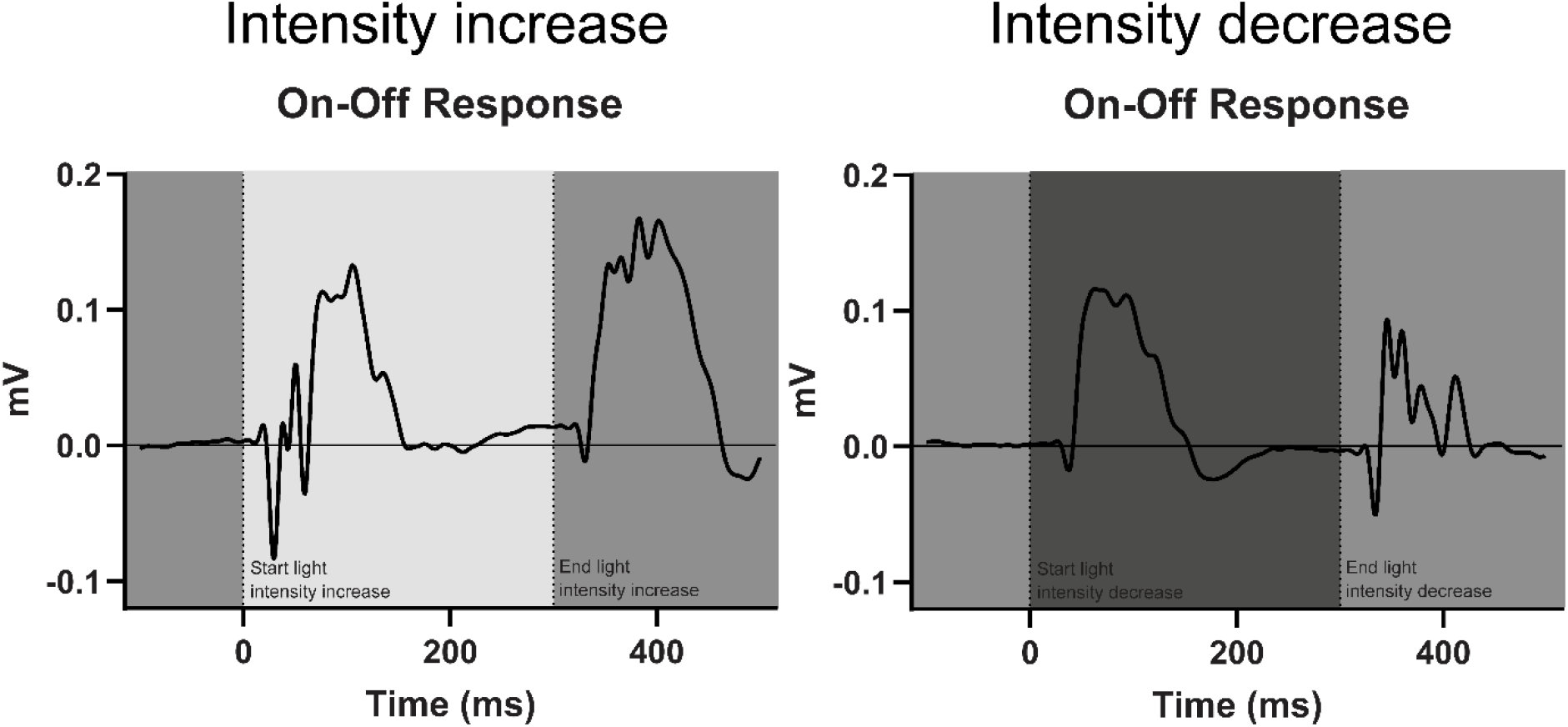
Comparison of VEP waveforms for the On- and Off-responses to a light intensity increase versus light intensity decrease. Light stimuli lasted for 300 ms, whereby these plots show the ‘On-response’ to the start of the 300-ms intensity increase or decrease as well as the VEP ‘Off-response’ to the light intensity changing back to baseline level. The On-response of the light increase, as well as the Off-response of the light decrease, concern a response to an increase in light intensity. The On-response of the light decrease, as well as the Off-response of the light increase, concern a response to a light intensity decrease. Presented data show the responses to a light increase and decrease, averaged over all mice for right and left V1 responses and 2 recordings on separate days.

